# Measurement of covalent bond formation in light-curing hydrogels predicts physical stability under flow

**DOI:** 10.1101/2024.06.30.601353

**Authors:** Jonathan M. Zatorski, Isabella L. Lee, Jennifer E. Ortiz-Cárdenas, Jeffrey F. Ellena, Rebecca R. Pompano

## Abstract

Photocrosslinking hydrogels are promising for tissue engineering and regenerative medicine, but challenges in reaction monitoring often leave their optimization subject to trial and error. The stability of crosslinked gels under fluid flow, as in the case of a microfluidic device, is particularly challenging to predict, both because of obstacles inherent to solid-state macromolecular analysis that prevent accurate chemical monitoring, and because stability is dependent on size of the patterned features. To solve both problems, we obtained ^1^H NMR spectra of cured hydrogels which were enzymatically degraded. This allowed us to take advantage of the high-resolution that solution NMR provides. This unique approach enabled the measurement of degree of crosslinking (DoC) and prediction of material stability under physiological fluid flow. We showed that NMR spectra of enzyme-digested gels successfully reported on DoC as a function of light exposure and wavelength within two classes of photocrosslinkable hydrogels: methacryloyl-modified gelatin and a composite of thiol-modified gelatin and norbornene-terminated polyethylene glycol. This approach revealed that a threshold DoC was required for patterned features in each material to become stable, and that smaller features required a higher DoC for stability. Finally, we demonstrated that DoC was predictive of the stability of architecturally complex features when photopatterning, underscoring the value of monitoring DoC when using light-reactive gels. We anticipate that the ability to quantify chemical crosslinks will accelerate the design of advanced hydrogel materials for structurally demanding applications such as photopatterning and bioprinting.

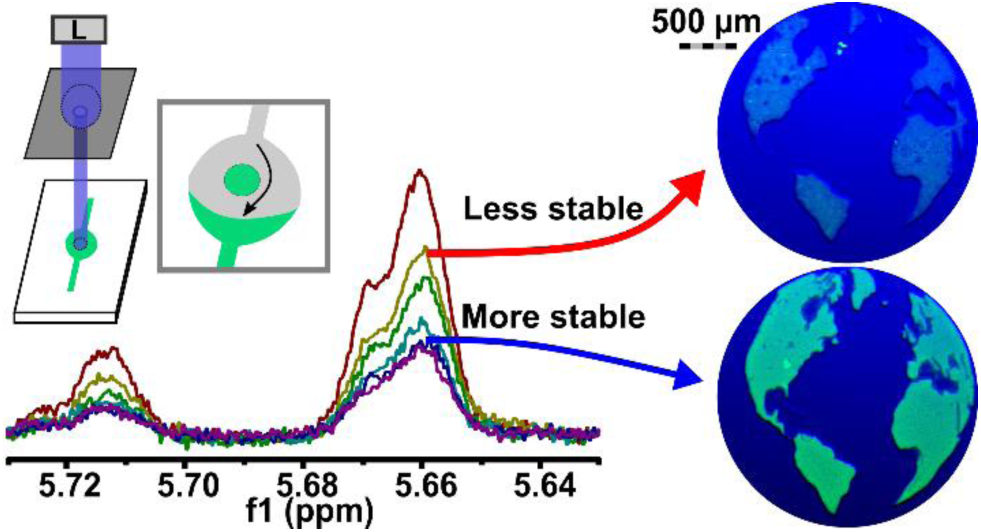

## INTRODUCTION

Nearly every branch of tissue engineering, from 3D cultured microphysiological models to implantable organs, demands well-controlled biomaterial chemistry. Modern applications require biomaterials that form and remain stable at specific locations and times, plus are tunable in terms of stiffness, porosity, and chemical content to meet the needs of each particular use case.^1,2^ These properties arise out of the chemistry of the base polymer as well as the chemistry used to crosslink it. Radical-induced polymerization, particularly initiated by light exposure, has found widespread use, because the properties of the biomaterial are readily tuned by adjusting the crosslinking conditions.^3–7^ However, many tissue engineering projects progress sluggishly through early optimization stages, striving to find reaction conditions that strike a balance between fabricating stable, high-resolution structures and the desired “soft-tissue” characteristics.^8^

Material stability in warm perfusion is an essential criterion for success in many microphysiological systems, bioreactors, and implants. Each of these contexts requires that the material maintain its integrity at physiological temperatures while immersed in culture media or body fluids, either of which may be flowing.^9^ Unfortunately, requirements for low stiffness in mimics of soft tissue can be in conflict with achieving stability under flow.^1,2^ This is particularly problematic when generating microscale patterned materials or other small features, which have high surface-to-volume ratio and decompose easily.^4^ To accelerate materials development in these contexts, a strategy for predicting the stability of a material based on its chemistry and reaction conditions is greatly needed.

Currently it is challenging to predict the physical stability of crosslinked materials, particularly under conditions of physiological perfusion. Rheometry is a standard approach, based on an underlying assumption that material stiffness is a proxy for stability.^10,11^ However, while stiffness has been linked to stability in air,^10^ it is often improperly interpreted as a proxy for stability in warm, submerged tissue engineering settings.^12^ Stiffness is impacted by several molecular parameters that may only indirectly impact stability, including rigidity, monomer length, phase transition, temperature, and solubility.^13,14^ Rather than stiffness, we reasoned that material dissolution may be more directly correlated with insufficient crosslinking.^15^ Therefore, we hypothesized that a method to assess the absolute degree of crosslinking (DoC) would be predictive of hydrogel stability.

Testing the relationship between stability and DoC requires advances in analytical measurement techniques, as current analytical methods struggle to report on this parameter. A development in rheology instrumentation enabled simultaneous stiffness and IR measurements during photocuring, but the low resolution afforded by IR prevented quantification of absolute DoC.^16^ Colorimetric assays based on consumption of functional groups are better suited for fractional rather than absolution quantitation,^17^ and furthermore require a unique detection reagent that is specific to functional group and not compromised by other gel components. Meanwhile, ^1^H NMR is well-suited for this analysis because the vinyl pendant groups often used for crosslinking reactions are clearly distinguishable from amino acid protons. Indeed, ^1^H NMR readily reported on degree of functionalization (DoF), which is the absolute content of photocrosslinkable groups such as methacryloyls or norbornenes (Scheme 1a).^17,18^ Yet, DoF only describes the upper limit of possible crosslinking, not the actual extent of crosslinks achieved under particular reaction conditions (Scheme 1b). A major obstacle for using ^1^H NMR to assess DoC is the decrease in spectral resolution that occurs as the material solidifies.^19–22^ Magic angle spinning solid-state (MAS) NMR partially addresses this issue, but still may not provide sufficient resolution for highly crosslinked gels.^19–22^ Therefore, precise measurement of DoC in crosslinked gels remains a significant challenge.

To fill these gaps, here we developed a method, utilizing ^1^H NMR and enzyme digestion, for assessing absolute DoC, with a focus on gelatin-based photocrosslinkable hydrogels, which are widely used to produce features of varied sizes in applications such as organs-on-chip, wound-healing applications, and 3D cell-culture,^4–7^ but suffer from unpredictable physical stability (Scheme 1c,d).^23,24^ After identifying that an enzyme digestion step successfully converted photopolymerized hydrogels to a state suitable for solution ^1^H NMR, we tested the extent to which DoC correlated with stability of two different materials under physiological conditions. We then characterized the relationship between stability and size, and finally tested the ability of the method to predict the stability of gel features having complex architectures.

**Scheme 1.**
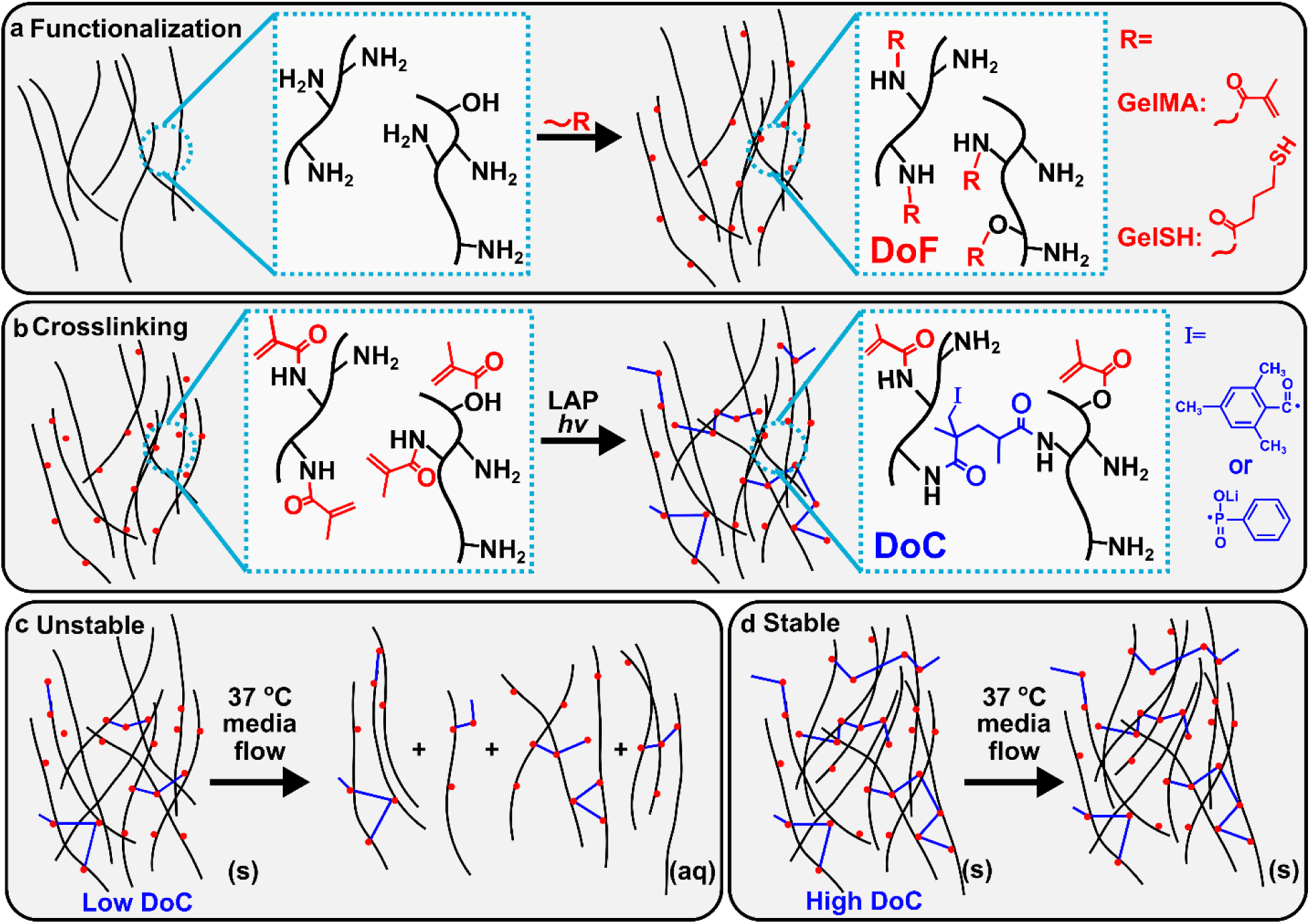
Natural ECM materials that are chemically functionalized and photocrosslinked may require a minimum crosslinking density to be stable. (a) Functionalization: Amino and hydroxyl terminated amino acids on gelatin (black lines) are functionalized with methacryloyl (gelMA) or thiol (gelSH) groups. (b) GelMA Crosslinking: Photopolymerization by blue or violet light and a photointiator such as lithium phenyl-2,4,6-trimethylbenzoylphosphinate (LAP) results in formation of covalent crosslinks and loss of methacryloyl vinyl groups. Bound radicals from the initiator are abbreviated as **I;** growing polymer chains could also bind at that location instead. (c) Instability: An insufficient degree of crosslinking (DoC) may result in material dissolution in the presence of warm media flow. (d) Stability: Sufficient crosslinking may result in a stable gel.

## RESULTS

### Digestion of hydrogel provided high-resolution ^1^H NMR spectra after photo-crosslinking

We and others have previously described ^1^H NMR methods to quantify the number of functional groups on gelatin, or degree of functionalization (DoF), based on the appearance of methacryloyl or norbornene groups (Fig. 1a).^17,18^ During crosslinking, those groups are consumed (Scheme 1b), which leads to the loss of signal in the vinylic range and potentially enables quantification of DoC.^19^ Indeed, by standard solution phase ^1^H NMR of the photocrosslinked material, we observed a decrease in peak height in the vinylic range when gelMA was exposed to 405 nm light for increasing lengths of time (Fig. 1b). However, we also observed peak broadening that pre-empted quantification, as expected due to loss of molecular motion and increase in nuclear reorientational correlation times within the crosslinked hydrogel.^25^ Therefore, we sought to develop an approach to obtain an ^1^H NMR spectrum from the crosslinked gels with sufficient resolution to quantify the vinylic methacryloyl peaks, similar to what was possible with the solution phase monomers.

**Figure 1.**
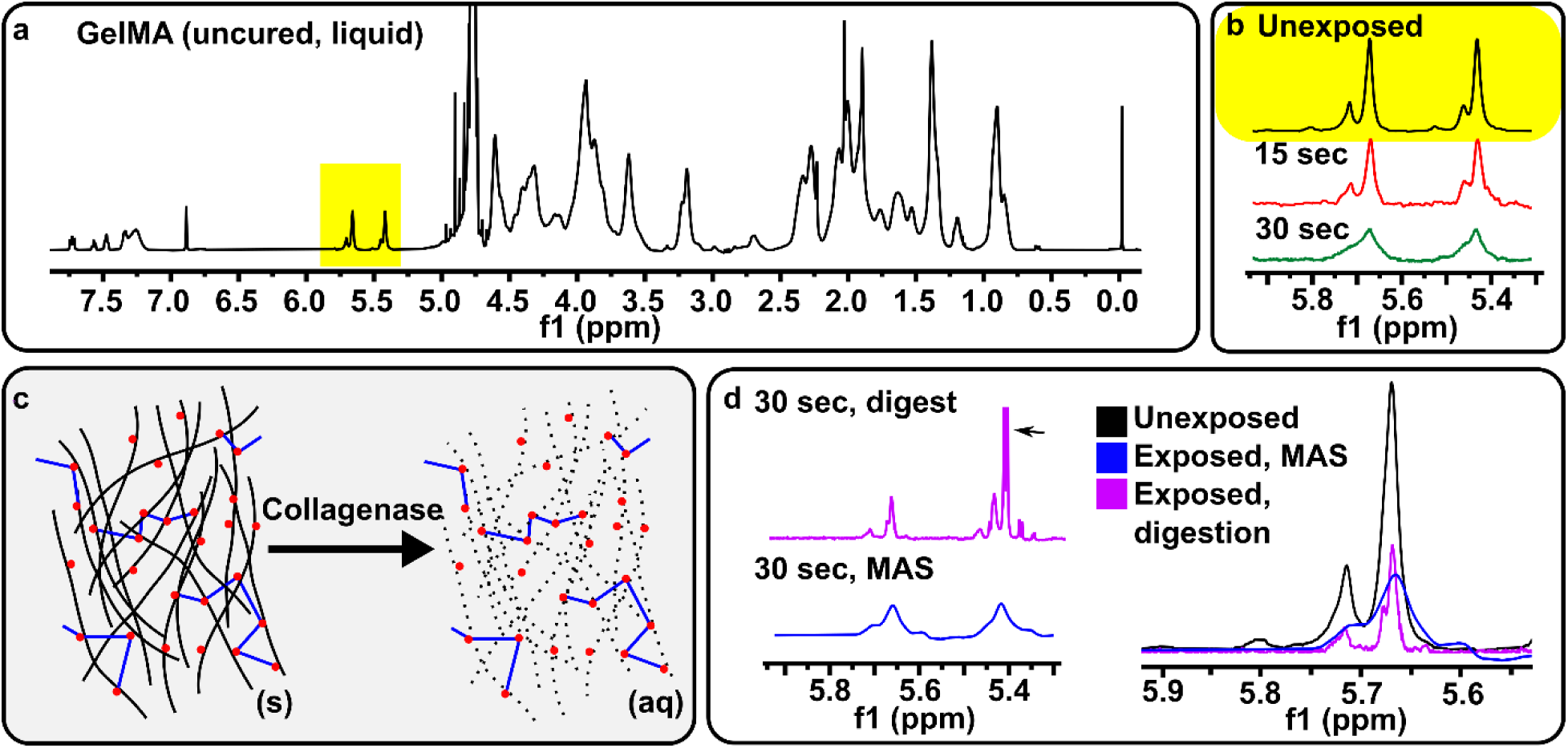
Digestion with collagenase D of gels produced high quality ^1^H NMR spectra. (a) Full solution phase ^1^H NMR spectrum of unexposed, 10% gelMA. Methacryloyl peaks highlighted in yellow. (b) Expanded view of H NMR spectra in the methacryloyl range for gelMA prior to exposure or after light exposure for 15 seconds (red) and 30 seconds (green), without enzymatic digestion. (c) Schematic illustrating the digestion of cured hydrogels: Collagenase D digested the gelatin backbone, leaving the synthetic crosslinks formed during photopolymerization in an aqueous solution. (d) H NMR spectra of exposed gel, analyzed either by MAS NMR (blue) or liquid state H NMR following enzyme digestion (magenta). Collagenase peak (arrow) interferes with upfield methacryloyl peak. Digestion provided higher resolution peaks in comparison to using MAS H NMR.

We reasoned that one could mitigate the peak line broadening due to increasing molecular size, while retaining the information on the number of functional groups, by selectively digesting the hydrogel at points other than the photocrosslinking site. The enzyme collagenase D cleaves at amides present in the gelatin peptide backbone, but does not cleave remaining methacryloyl pendant groups (Fig 1c).^26^ Therefore, we tested the extent to which collagenase digestion would convert the solid cured gel into a liquid without affecting the analysis. To circumvent the need for solvent exchange prior to analysis (H_2_O to D_2_O), we digested the hydrogel in D_2_O solvent. The hydrogel digested as expected, as collagenase activity is similar in aqueous and deuterated solvents.^27^ Analysis of the digested solution by standard solution ^1^H NMR showed well resolved peaks of lower intensity than from the unexposed gelMA precursor, consistent with loss of the vinyl group during photocrosslinking (Fig 1d). By obtaining a ^1^H NMR spectrum of collagenase (Fig. S1), we identified the sharp peak at 5.40 ppm in the digested sample as interference from collagenase. We compared this method to the gold standard for analysis of solids, magic angle spinning (MAS) H NMR. Digestion of the gel yielded a sharper peak than MAS NMR, and in fact the line width was narrower than the unexposed gelMA precursor (Fig 1d). We attribute the higher resolution to the lower average molecular weight caused by digestion.

### Enzyme digestion enabled reaction monitoring of gelMA photocrosslinking

Having shown that digesting an exposed gel improved spectral resolution, we investigated whether digestion would enable monitoring the photocrosslinking reaction of gelMA. To calculate crosslinking density (DoC) for gelMA, we reasoned that we could quantify the loss of apparent DoF to calculate the number of crosslinks formed. We first measured the apparent DoF of the sample by comparing the integral of the downfield methacryloyl peak (5.60 to 5.75 ppm) to the integral of the internal standard (Eq. 1), using sodium trimethylsilylpropanesulfonate (DSS) as an internal standard, as previously reported.^17^

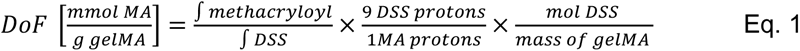

DoC was estimated at each timepoint by subtracting the apparent DoF at time *n* (*t_n_*) from the initial DoF at time zero (*t_0_*), then dividing by a factor equal to the number of functional groups consumed per crosslink (*f)* (Eq. 2). For gelMA, we assumed *f*=2, accounting for two methacryloyl groups per gelMA linkage.

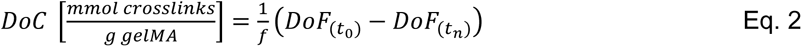

Similarly, we reasoned that the maximum possible DoC would be reached once every functional group was consumed, which for gelMA would occur if every methacryloyl group reacted with one other (Eq. 3).

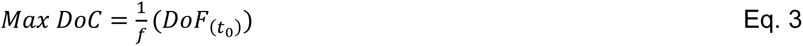

We note that the use of *f*=2 for gelMA provides a lower limit of the max DoC, as some crosslinks may include three or more methacrylates due to the chain growth mechanism. We reasoned that the distance between methacrylate functionalizations on the protein would favor dimeric crosslinks over longer chains.

To test the effectiveness of this approach for calculating DoC, we photo-exposed gelMA in five second increments and analyzed the resulting hydrogels after enzymatic digestion. The initiator, LAP, has an absorbance maximum at 375 nm, so faster reactions were expected at 385 nm than 405 nm.^28^ We observed a clear decrease in peak area with exposure time at both wavelengths, consistent with a loss of methacryloyl vinyl groups due to covalent bond formation (Fig 2a,b). Converting the peak area to DoC vs time resulted in a sigmoidal growth curve upon 405 nm exposure, and an accelerated growth curve upon 385 nm exposure, as expected (Fig. 2a-c). Interestingly, both conditions levelled off at 70 % of the predicted Max DoC, suggesting that steric hindrance may have prevented all the methacryloyl groups from reacting.^19,29^

**Figure 2.**
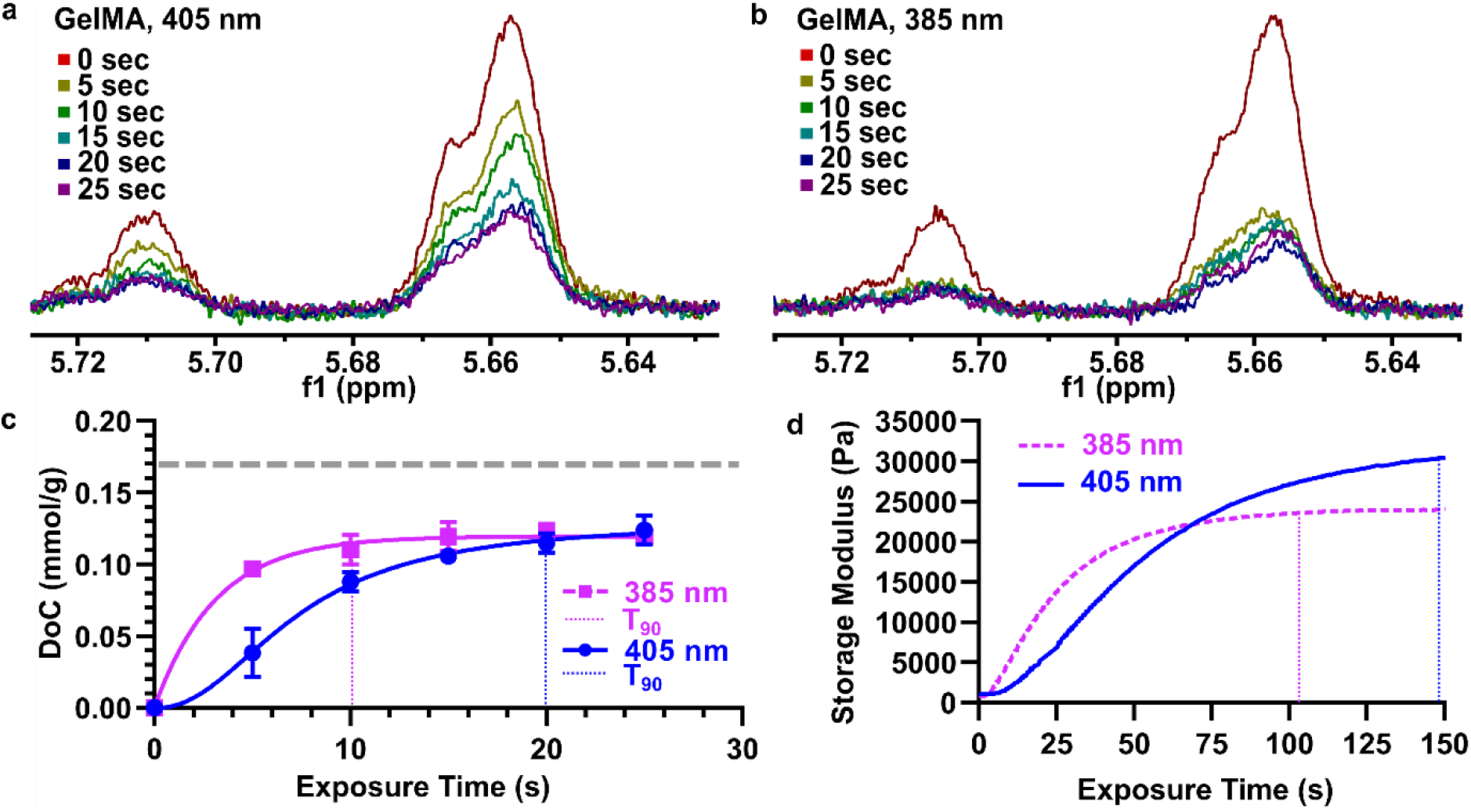
Quantification of degree of crosslinking (DoC) in gelMA using collagenase D prior to liquid state H NMR. (a) H NMR spectra of 10% gelMA exposed to 405 nm, 25 mW cm^-2^ light. Digestion with enzyme followed by H NMR enables tracking of DoC with 5 second exposure intervals. (b) H NMR spectra of gelMA exposed to 385 nm, 25 mW cm^-2^ light. (c) Quantification of DoC from spectra shown in Fig. 2a,b. Plot of 10% gelMA DoC [mmol/g] versus exposure time with 405 nm and 385 nm light. Max DoC represented by grey dotted line. n=3, points represent mean, and error bars represent standard dev. 405 nm and 385 nm curves fit to sigmoidal and exponential functions, respectively. (d) Time-dependent rheological characterization of gelMA. Plot of gelMA storage modulus [Pa] versus exposure time with 405 nm and 385 nm light, 25 mW/cm^2^ intensity.

Finally, as stiffness is often used as a proxy for extent of crosslinking, we tested the extent to which stiffness correlated with DoC as the reaction progressed. While the DoC plateaued within the first 10 – 20 seconds (t_90_ = 20.0 sec at 405 nm; t_90_ = 10.8 sec at 385 nm), whereas the stiffness plateaued at least 90 seconds later (t_90_ = 147 sec at 405 nm; t_90_ =103 sec at 385 nm), for both wavelengths of light (Fig. 2d). Thus, the stiffness increased beyond the time when the methacryloyl peaks disappeared. These results may indicate that methacryloyl groups were consumed by radical reactions more quickly than covalent crosslinks formed, or may indicate the continuing of a physical gelation process after crosslinking is complete.

### Stability of gelMA features was related sigmoidally to DoC

Now that we could quantify DoC, we tested its relationship with the physical stability of photocured gels. More crosslinking is generally associated with greater stability, and we hypothesized that the relationship could be either linear or sigmoidal.^15^ To test this hypothesis, we challenged photocrosslinked regions of gel with conditions of steady-state flow at physiological temperature, as is found in many organ-on-chip experiments.

Freestanding islands of gel were photopatterned inside a simple microfluidic flow cell (Fig 3a), following procedures previously described by us and others.^4,30^ The chip was filled with fluorescently labeled polymer precursor solution, a photomask laid over the chip, and the precursor exposed through a 800 µm diameter pinhole. This simple photopatterning strategy made it straightforward to evaluate multiple exposure times on a single chip, by exposing one region, re-positioning the mask, re-exposing, and so on (Fig. 3b). To observe how stability trended with changes in DoC, we tested timepoints preceding the plateau (T_90_) in DoC vs time (Fig. 2c). After exposure of four regions (e.g. for 2, 4, 6, and 8 seconds) for local gelation, the remaining uncrosslinked precursor was rinsed away. Finally, the chips were perfused with warm PBS for two hours.

**Figure 3.**
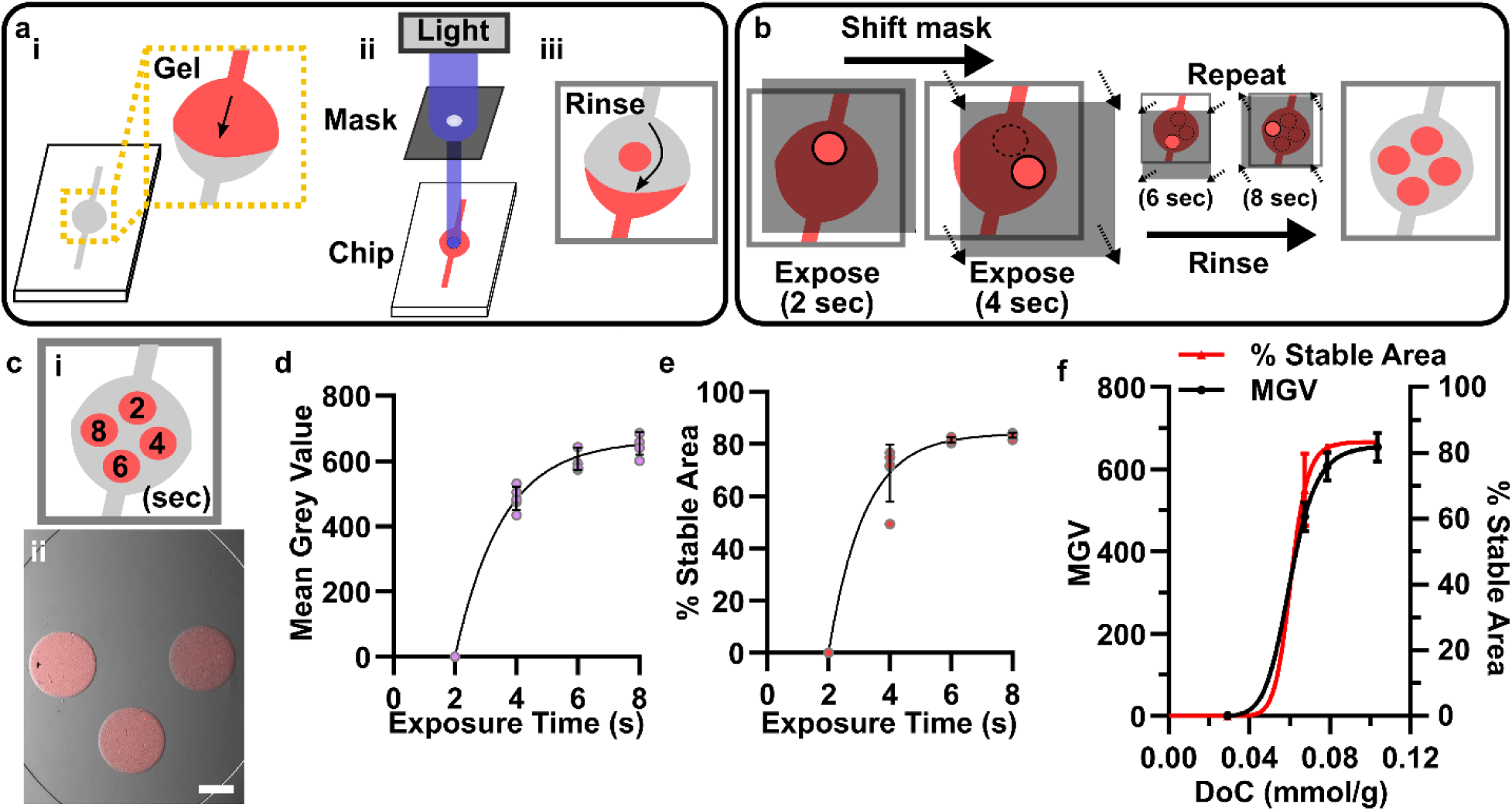
Stability of patterned GelMA under physiological fluid flow had a threshold dependence on DoC. (a) Schematic of *in situ* photopatterning process.^4^ (i) Chip filled with gel precursor. (ii) Photomask aligned with chip, clamped (not shown for clarity), and exposed. (iii) Unexposed gel precursor rinsed away with PBS. (b) Schematic of patterning with multiple exposure times on a single chip by shifting the photomask and performing sequential exposures. (c) Microphotograph of circular features of 10% gelMA-rhodamine (red) after varied exposure times with 385 nm, 25 mW cm^-2^ light. (i) Schematic of the position of each exposure time point. (ii) Composite (fluorescence overlayed over brightfield) microscope image of the chip following a PBS rinse (10 min, 5 µL/min, 50 µL) and physiological fluid flow (2 hours, 5 µL/min, 600 µL). Scale bar = 500 µm. (d) Quantification of fluorescence intensity (measured in mean grey value) of circular gelMA features from Fig. 3c. (e) Quantification of % stable area of circles, defined as the area of the patterned gel divided by the area of feature in the photomask. Data were fit to an exponential curve. (f) Plot of mean grey value and % stability versus DoC (mmol/g) of gelMA features. Data were fit to a sigmoidal curve. n=5 chips.

Under these conditions, both fluorescent intensity as well as the percent stable area of the gel increased with exposure time (Fig. 3c-e). GelMA areas that were photo-exposed for at least 6 seconds were stable, while areas exposed for 2 seconds were nearly always unstable and washed away completely (Fig. 3c). The 4-second exposure was semi-stable, as some fraction of the pattern was always present after rinsing, but it was never complete. To account for the increased shear stress on patterns closer to the inlet of the flow cell, we rotated the exposure order and observed no difference in stability.

When plotted against DoC obtained from NMR measurements at the same exposure times (from Figure 2), both fluorescent intensity and % stable area were well fit by a sigmoidal curve, with a DoC-at-half-max of 0.060 mmol/g (Fig 3f). Thus, physical stability of the photopatterned gelMA features appear to require that the crosslinking density surpass a critical DoC threshold (Fig 3f). Because gel stiffness also increased with exposure time (Fig 2d), we cannot exclude that changes in stiffness may have driven this behavior. However, because stiffness increased over a 10-fold longer period of exposure (at 385 nm, t_90_ = 103 sec for stiffness; t_90_ = 10.8 sec for DoC), we consider it more likely that DoC was responsible for the dramatic increase in stability between 2 and 8 sec than stiffness.

### Stability of features in a gel with a step-growth crosslinking mechanism was related exponentially to DoC

To determine the extent to which the sigmoidal relationship between DoC and stability was generalizable from GelMA to other crosslinking mechanisms, we assessed the same metrics using a semi-synthetic polymer blend composed of thiol-modified gelatin and a norbornene-functionalized polyethylene glycol (PEG) linker (gelSH-pegNB) (Fig. 4a). Unlike gelMA, which crosslinks via a chain-growth mechanism, gelSH-pegNB crosslinks via step-growth mechanism.^7^ Thiol-norbornene hydrogels have seen widespread use due to increased compatibility with cell culture and *in situ* photopatterning.^4,7,31^

**Figure 4.**
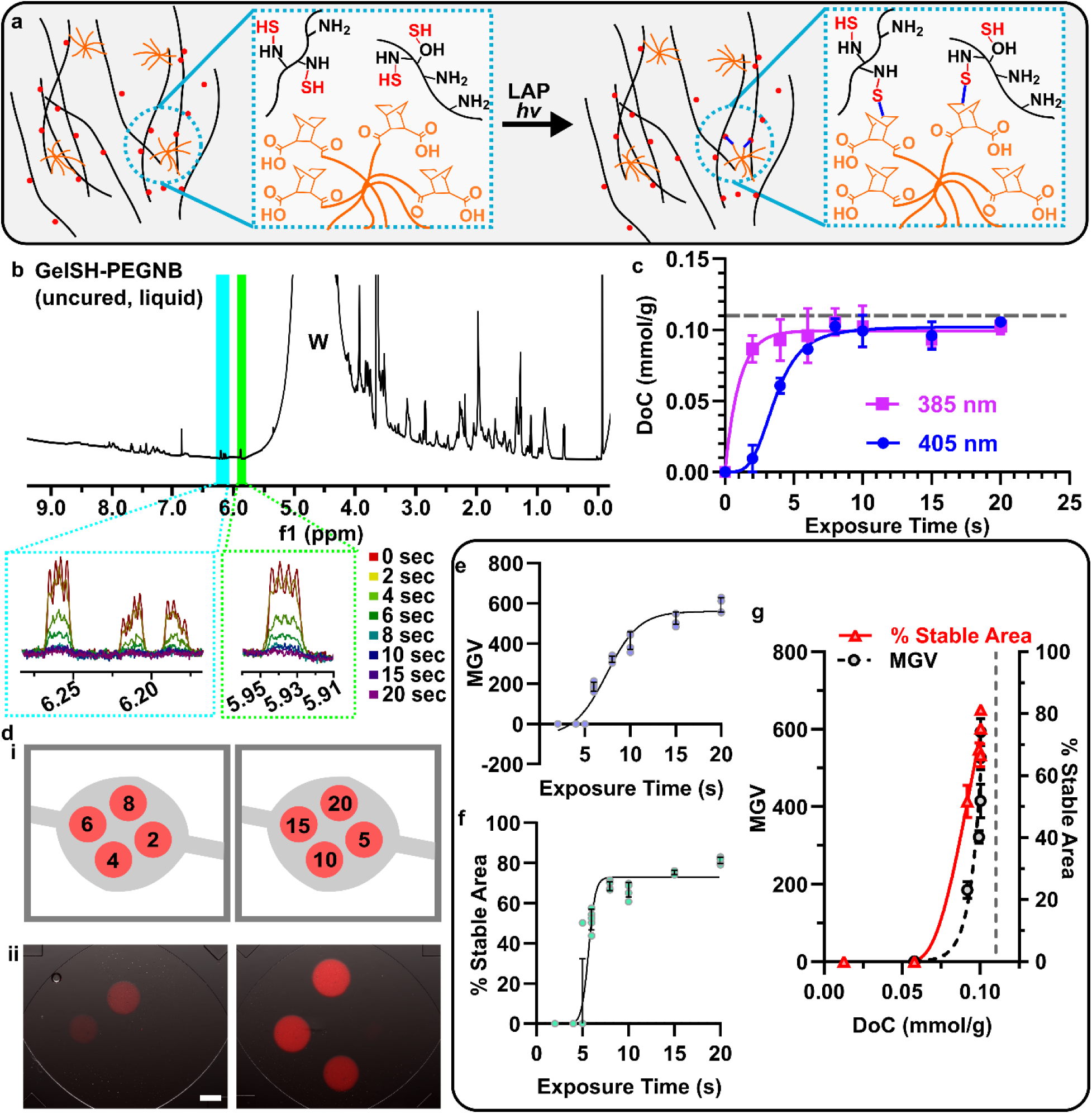
Stability of a step-growth hydrogel had a threshold dependence on DoC. (a) Schematic illustrating the crosslinking reaction between gelSH and 8-arm PEG-NB. (b) Full H NMR spectrum of uncured, unexposed gelSH-pegNB. NB peaks highlighted in blue and green. Zoomed in boxes show norbornene peaks after various exposure times. Water peak identified as “W.” (c) Quantification of DoC versus exposure time with 405 nm and 385 nm light. Grey dotted line indicates DoC_max_. Each point represents the mean from n=3, error bars represent standard deviation. (d) Circular features of gelSH-rhodamine features exposed for differing times with 405 nm light. (i) Position of each exposure time point. (ii) Composite (fluorescence overlaid over brightfield) microscope image of chip following PBS rinse (10 mins, 5 µL/min, 50 µL) and physiological fluid flow (2 hours, 5 µL/min, 600 µL). Scale bar = 500 µm. (e) Quantification of fluorescence intensity (mean grey value) of circular gelSH features from Fig 4d. (f) Quantification of percent stable area (area of circular feature/area of circular photomask) of circular gelSH features from Fig 4d. Data fit to a sigmoidal curve. (g) Plot of MGV and % stable area versus DoC (mmol/g) of gelSH features. Data fit to an exponential curve. Grey dotted line indicates DoC_max_. n= 5 chips. Each point is a pattern from a chip. Error bars represent standard deviation.

Using enzymatic digestion prior to H NMR, a decrease in peak area of the vinyl norbornene protons was easily observable upon photocrosslinking (Fig. 4b).^17,31^ DoF of gelSH-pegNB was calculated using a ratio of NB protons per DSS protons is 2:9 (Eq. 4):

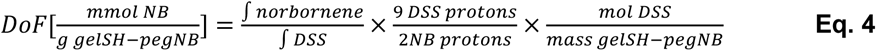

We calculated DoC as described above (Eq. 3), using *f*=1 to represent that each crosslink consumed a single norbornene group. Interestingly, unlike gelMA, the gelSH-pegNB hydrogel approached the DoC max, with peak-heights of vinylic protons meeting or nearly meeting the baseline at exposures beyond 8 sec and the DoC curve leveling off at 91% of the theoretical maximum (Fig. 4b,c).

To determine how DoC was related to stability for this material, we performed *in situ* photopatterning with gelSH-pegNB using the same approach as above (405 nm; Fig. 4d). Both the fluorescence intensity and percent stable area increased sigmoidally with exposure time, with the latter increasing faster (Fig. 4e,f). Thus, for both gels, stability under fluid flow required a DoC above a critical minimum value. However, unlike gelMA, where fluorescence intensity and stable area leveled off as DoC increased, for gelSH-pegNB they continued to increase exponentially even as DoC leveled off near its theoretical maximum (Fig. 4g). This result is strikingly reminiscent of well-established polymer theory (Fig. S2), which states that chain-growth polymers, such as gelMA, achieve maximal molecular weights in relatively fewer crosslinking steps than step-growth polymers, such as gelSH-pegNB.^32,33^ We posit that the stability of patterned hydrogels is related to the extent of the crosslinked network and thus to molecular weight. Such a relationship is an interesting direction for future work.

### Higher DoC was required to stabilize smaller features

A major advantage of photopatterning is the ability to produce different sizes of features on demand, but smaller patterned features are often more susceptible to erosion from fluid flow than larger ones.^4^ Although higher DoC has been suggested to improve the stability and resolution of relatively smaller features when bioprinting, it must be not be too high or one risks lowering permeability and cell viability due to low porosity.^34^ Therefore, the extent of crosslinking must be optimized for each feature size. We hypothesized that smaller features may have a higher threshold DoC for stability.

To test this hypothesis, we photopatterned gelSH-pegNB through a size-array photomask with varied exposure times, followed by perfusion for two hours with warm PBS. The mask was comprised of three circles that were 200, 400, and 600 µm diameter (Fig 5a), in triplicate. These results were combined with the 800 µm diameter data from Fig 4d. As predicted, percent stable area was clearly dependent on feature size, both at individual timepoints (Fig. 5b) and in the threshold DoC required for stability (Fig. 5c). We arbitrarily chose 35% stable area to quantify the threshold (DoC_35%_), and found that the DoC_35%_ increased for smaller feature sizes (DoC_35%_ = 0.099 at 400 µm; 0.091 at 600 µm; 0.086 at 800 µm, in units of mmol-crosslinks/g gel) (Fig. 5c). 200 µm features were not stable at any exposure time. Interestingly, unlike % stable area, the fluorescence intensity of any features that did form was largely independent of feature size (Fig. 5d,e). Thus, a higher number of crosslinks were required for smaller, free-standing features to resist erosion at the edges due to fluid flow, but the height and local concentration of gel remaining after erosion was independent of feature size. These data are consistent with the uniform light exposure during crosslinking producing a uniform DoC across all features regardless of their size, with loss of stability at the edges due to diffusive loss of uncrosslinked monomers from the hydrogel.

**Figure 5.**
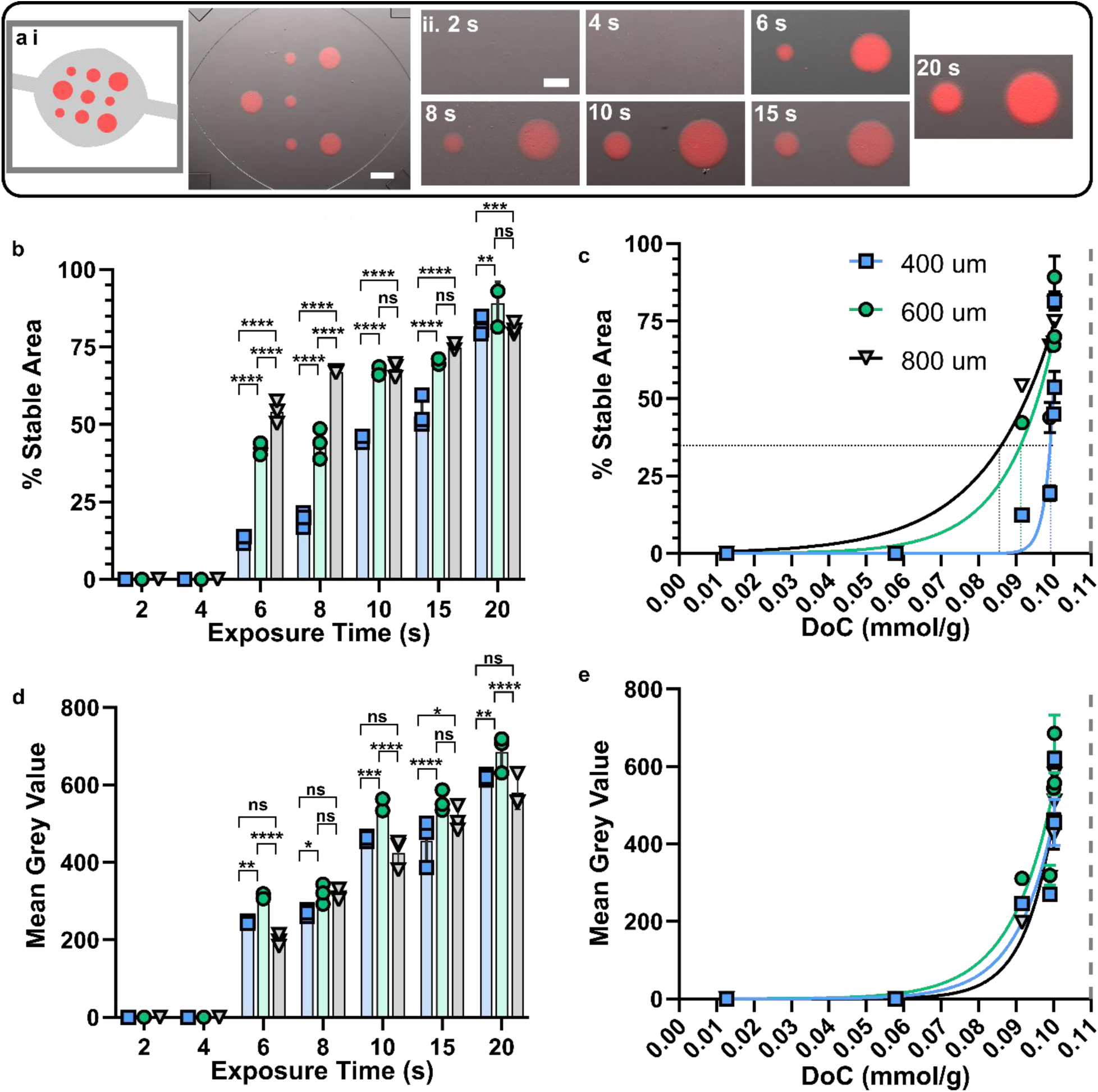
The DoC threshold for the stability of photopatterned features was inversely proportional to size. (a) (i) Schematic and (ii) Image of 5% gelSH with 20 kDa 8-arm PEG-NB photopatterned as an array of 200, 400, and 600 µm circles with 405 nm light 25 mW cm^-2^ intensity at varying time points. Scale bar = 500 µm. (iii) Zoom on 400 and 600 µm patterns for each timepoint. Scale bar = 250 µm. (b,d) Quantification of % stable area (b; area of circular feature/area of circular photomask) and fluorescence intensity (d; mean grey value) of circular gelSH features from Fig. 4d and 5a. Bars show mean and std dev; n = 3. Ordinary one-way ANOVA. (c, e) Plot of % stability (c) and mean grey value (e) versus DoC (mmol/g) of gelSH features. Dots show mean and standard deviation. 35% stability represented by black, green, and light blue vertical dotted lines. Data fit to an exponential curve. Max DoC represented by grey dashed line. n= 3 chips.

### The stability of complex photopatterned features followed expected dependence on DoC

Finally, we applied this concept of a size-dependent critical DoC to predict the cross-linking conditions required for a complex pattern with features of various size and connectivity: the Atlantic face of Earth (Fig. 6a-e). We hypothesized that the relationship between critical DoC and feature size (Fig. 5c) would hold for the various continents and island features. For example, we predicted larger islands such as Greenland (120 µm mean radius) and South America (322 µm mean radius) would be stable at a lower threshold (DoC_35%_) than a smaller island such as Panama (49 µm mean radius).

**Figure 6.**
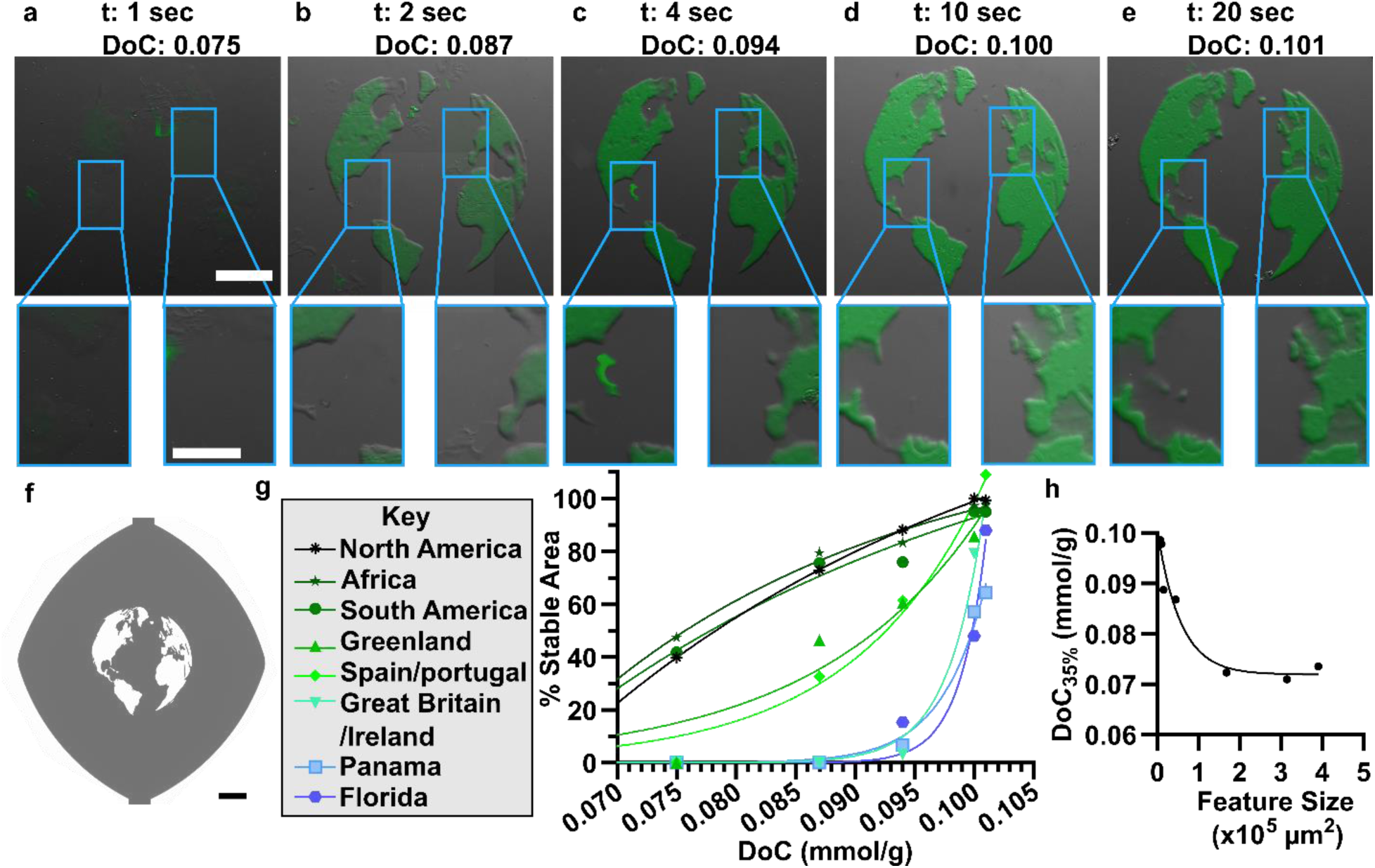
The stability of complex features was predicted using DoC and size. GelSH-pegNB was exposed with 385 nm light 25 mW cm^-2^ intensity through a photomask of the Earth. (a-e) Composite fluorescence microscopy images of gelSH-pegNB features following exposure through an Earth photomask at various expo sure times. The corresponding DoC values (± 0.01 mmol bonds/g gel) are shown in the top right of each image. Scale bar is 500 µm. Blue squares show zoomed in views of island features, with scale bar of 250 µm. (f) Photomask for generating the land features from the Earth’s Atlantic face. Scale bar= 500 µm. (g) Plot of DoC vs percent stable area for Earth continents and islands, fitted with exponential growth or plateau curves. (h) Plot of DoC_35%_ vs feature area, determined from exponential curves shown in (f).

To test these predictions, the Earth pattern was photopatterned in the microfluidic flow cell and perfused for two hours as in prior experiments. To generate features below 200 µm diameter, we accelerated crosslink formation by exposing with 385 nm light (Fig. 4c). This is unlike figure 5, in which gels were exposed to 405 nm light. Perfusion ran from the north to south poles. As expected, larger features were more stable at lower exposure times (Fig. 6a-e; Table 1). We saw that the larger continents (Africa, North and South America) were crosslinked sufficiently to resist flow at 0.07 mmol crosslinks/g gel (Fig. 6b). Slightly smaller features like Greenland and Spain became stable at 0.087 and 0.089 mmol crosslinks/g gel, respectively (Fig. 6c). Isle nation features like Great Britain and the Caribbean islands, as well as land that directly obstructed flow paths, such as Panama, were not sufficiently crosslinked for stability until at least 0.098 mmol crosslinks/g gel were formed (Fig. 6d,e). Features up to 150,000 µm^2^ were best fit to an exponential growth curve, and features above 150,000 µm^2^ were best fit to an exponential plateau curve (Fig. 6f). These differing curves may both ultimately reflect a sigmoidal dependence; we were not able to apply < 1 sec exposure times and thus could not access lower DoC.

**Table 1.** Earth photomask feature dimensions and experimentally measured critical DoC.

| Feature | Area ($\mu\text{m}^2$) | Mean radius ($\mu\text{m}$) | Edge/Area | DoC<sub>35%</sub> [mmol crosslinks/ g gel] |
| --- | --- | --- | --- | --- |
| Earth | 3,012,172 | 979.2 | N/A | N/A |
| Africa | 314,658 | 316.5 | 0.010 | 0.071 |
| North America | 390,503 | 352.6 | 0.014 | 0.074 |
| South America | 168,463 | 321.6 | 0.012 | 0.072 |
| Greenland | 45,136 | 119.9 | 0.024 | 0.087 |
| Spain/Portugal | 15,559 | 70.4 | 0.048 | 0.089 |
| Great Britain/Ireland | 10,563 | 58.0 | 0.047 | 0.098 |
| Panama | 7,418 | 48.6 | 0.078 | 0.099 |
| Dominican Republic/Haiti | 1,361 | 20.8 | 0.112 | N/A<sup>1</sup> |

<sup>1</sup> Not able to calculate DoC<sub>35%</sub>, because feature was only detectable at the highest tested DoC.

Intriguingly, using the exponential fits from Fig 6g, we observed an exponential decay relationship between feature area and the threshold DoC_35%_ for stability (Fig. 6h). As diffusivity within a hydrogel in general is inversely proportional to DoC,^35,36^ we speculate that this result may suggest a dependence on diffusive loss of uncrosslinked polymer for instability, reminiscent of other diffusion-driven, swelling-induced instabilities of hydrogels.^37,38^

In summary, the hypothesis that smaller features in a complex pattern would require a higher DoC to be stable was supported in the Earth pattern, and in fact the critical DoC decreased exponentially with feature area.

## CONCLUSION

In summary, here we used solution NMR to directly quantify the absolute crosslinking density within photocrosslinking hydrogels for the first time. Solution NMR of collagenase digested, crosslinked gels yielded higher resolution spectra than MAS NMR of undigested gels. This measurement strategy revealed that the stability of photopatterned gelMA and gelSH-pegNB had a threshold dependency on DoC, and that smaller photopatterned features required higher DoC to resist erosion under perfusion. Finally, we found that the critical DoC depended linearly on feature area in complex features, which may enable explicit prediction of photopatterning conditions as a function of feature size in future work. Thus, we anticipate that DoC quantification will enable advanced biofabrication of soft materials to meet architecturally demanding designs, such as photolithography and bioprinting.

## METHODS

### Hydrogel material and preparation

Hydrogels were prepared as described previously.^4^ Briefly, precursor solution for gelMA was prepared by combining 60% functionalized gelatin methacryloyl (bloom 300, Sigma Aldrich Lot: MKCQ6360) and the photoinitiator, lithium phenyl-2,4,6-trimethylbenzoylphosphinate (LAP, Sigma Aldrich), to a final concentration of 10% w/v gel and 0.01% w/v LAP. Precursor solution for gelSH-pegNB was prepared by combining thiol-modified gelatin (Sigma Aldrich, Lot: MKCJ5413, 0.223 mmol –SH/g gelatin) with 8-arm PEG-NB 20 kDa (Creative PegWorks) and LAP for a final concentration of 5% w/v gelSH, 10 mM PEG-NB and 0.01% w/v LAP.

For H NMR analysis, the internal standard, sodium trimethylsilylpropanesulfonate (DSS, Sigma Aldrich) was added at 0.15 mM, and the precursor was prepared in D_2_O. For *in situ* photopatterning and rheometry, the precursor was prepared in 1x phosphate buffered saline with no calcium and no magnesium (PBS, Gibco, item # 14190250). After combining ingredients, the precursor solution was incubated at 40 °C for two hours and used within one day. Precursor was stored at 37 °C for one hour prior to use, or otherwise stored at 4 °C. All other chemicals were provided by Thermofisher.

### Photopolymerization and enzyme digestion

Precursor solution was polymerized in a 12 well plate by exposing for various times at 405 or 385 nm using collimated LEDs at an intensity of 25 mW/cm^2^. Specifically, a Prizmatix LED (UHP-F-405 or UHP-F-385) fitted with a liquid light guide and 90° reflector attachment with collimating optics was aligned to expose an area the size of a typical 12.7x8.5 cm microwell plate.

Following exposure, 0.3 mL of a solution of 0.7 mg/mL collagenase D and 5.5 mg/mL CaCl_2_ (Sigma Aldrich) in D_2_O was added to each well and mixed with a pipette tip, then incubated at 37 C for at least 12 hr to generated a digested solution.

### NMR analysis

Solution ^1^H NMR was performed on a Bruker Avance III 800 MHz NMR spectrometer with helium cooled cryoprobe. Total volume of solution was 600 uL. 8 and 32 scans were obtained for ^1^H spectra of methacrylates and norbornenes, respectively. Hydrogel samples that were analyzed without digestion were photoexposed *in situ* within the transparent, glass NMR sample tube.

For solid-state NMR analysis, precursor solution was added to the rotor, and the uncapped rotor was placed upright underneath the LED light path. After photoexposure, the rotor was capped and spectra was collected immediately. ^1^H spectra were obtained using a 500 MHz Varian VNMRS NMR spectrometer with 3.2 mm T3 HXY MAS probe. Single 90° excitation pulses (2.2 µs) were used and the time between pulses was 12 sec. Spin rate was 6000 Hz.

Mestrenova (v14.3.0-30573) was used to analyze all spectra. Baseline correction was performed using polynomial correction. Zero (ph0) and first-order (ph1) phase correction was used to minimize phase error. The internal standard peak was identified and set to 0 ppm. Methacryloyl peaks were identified and integrated at 5.62 to 5.72 ppm. Norbornene peaks were integrated at 5.92 to 5.95 ppm and 6.07 to 6.27 ppm and added together to give final norbornene peak area.

### Rheological characterization

Rheological characterization was performed on a MCT302 Anton Parr Rheometer. Thirty µL of precursor solution was pipetted onto a light-transmitting stage, and the Prizmatix LEDs (described above) were positioned underneath the stage. Hydrogel storage modulus was measured by equipping the stage with a 20 mm parallel plate and using time sweep mode at 5% strain with a 0.1 mm gap and 1 Hz frequency. Baseline shear storage modulus was measured, followed by measurement during irradiation at 25 mW/cm^2^.

### *In situ* photopatterning

*In situ* photopatterning was performed exactly as described in our previous work,^4^ with one key exception. When assessing pattern stability by shifting the photomask on chip to test multiple timepoints, the photomask was shifted prior to each sequential exposure, then uncrosslinked gel was rinsed out after all four patterns were produced. Photomasks for preparing master molds and *in situ* photopatterning were drawn in AutoCAD LT 2019 and printed by ArtNet Pro Inc at 20,000 DPI.

### Microscopy

Microscopic imaging was performed using an upright Zeiss AxioZoom microscope equipped with an Axiocam 506 mono camera, HXP 200C metal halide lamp, and PlanNeoFluor Z 1× objective (0.25 NA, FWD 56 mm). Fluorescence of rhodamine-labeled materials was detected using Zeiss Filter Set 43 HE (Ex: 550/25, Em: 605/70). Brightfield images were collected using transmitted light from a ZEISS Cold Light Source CL 9000 LED. Zen Blue software was used for image collection.

### Image analysis

Images were analyzed in ImageJ v1.53t. To detect NHS rhodamine-labeled patterned features, a threshold was set and 16-bit fluorescence images were converted to binary. The particle detection tool was used to define regions and quantify their mean fluorescence and size. % fractional area was determined by dividing the measured area of the pattern by the area of the pattern in the photomask.

### Statistical analysis

Data was prepared and statistical analysis was performed using GraphPad Prism 8.4.2.

## AUTHOR CONTRIBUTIONS

JMZ, ILL, and RRP designed the study, assessed the findings, and drafted the manuscript. JMZ and ILL performed experiments. JFE guided NMR optimization and acquired solid-state NMR spectra. JOC contributed preliminary findings and interpretations that informed the design of the study.

## ACKNOWLEDGEMENTS

This work was supported by the National Institute of Biomedical Imaging and Bioengineering (NIBIB) under Award Number U01EB029127 through the National Institutes of Health (NIH), with co-funding from the National Center for Advancing Translational Sciences (NCATS). JMZ was supported in part by the National Science Foundation Graduate Research Fellowship Program (NSF GRFP). JMZ and IIL were supported in part by the Double Hoo Research Grant from the University of Virginia. IIL was supported in part through the Lester Andrews Undergraduate Research Fellowship from the Department of Chemistry at the University of Virginia. This work utilized the UVa Biomolecular Magnetic Resonance Facility, which is supported by the University of Virginia School of Medicine, Research Resource Identifiers (RRID): SCR_018736. The authors thank Dr. Steven Caliari for graciously allowing us to perform rheological characterization of gelMA within his space. The authors also thank Jan-Willem Lamberink-Ilupeju, PhD (Western University) for helpful discussions about gelMA crosslinking mechanisms.

## SUPPLEMENTARY DATA

Supporting information (Figures S1 and S2) are available in the SI document. Additionally, representative source data generated in this study are posted under Zatorski et al., "Replication Data for: Measurement of covalent bond formation in light-curing hydrogels predicts physical stability under flow", https://doi.org/10.18130/V3/BSVZXH, University of Virginia Dataverse.

