## Supporting Information for "Measurement of covalent bond formation in light-curing hydrogels predicts physical stability under flow"

Electronic Supplementary Materials for

**Contents:**


- Supporting Figure S1: Collegenase D $^1$H NMR spectrum
- Supporting Figure S2: The stability of GelMA and GelSH-pegNB trends with DoC in a manner that is analogous to polymer theory.
- Supplemental references.

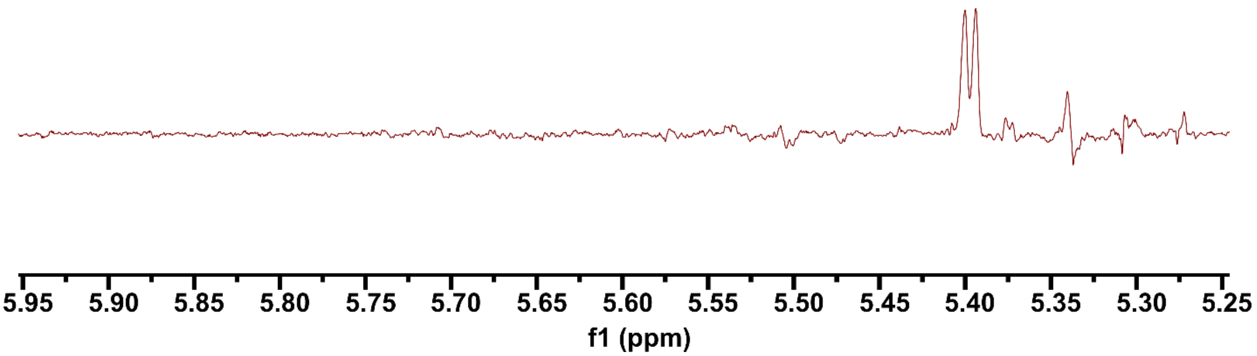


**Supporting Figure S1.** Zoom of $^{1}H$ NMR spectrum of collagenase D from *Clostridium histolyticum*, dissolved in $D_2O$ at 1 mg/mL (peaks at 5.4 ppm). 64 scans.

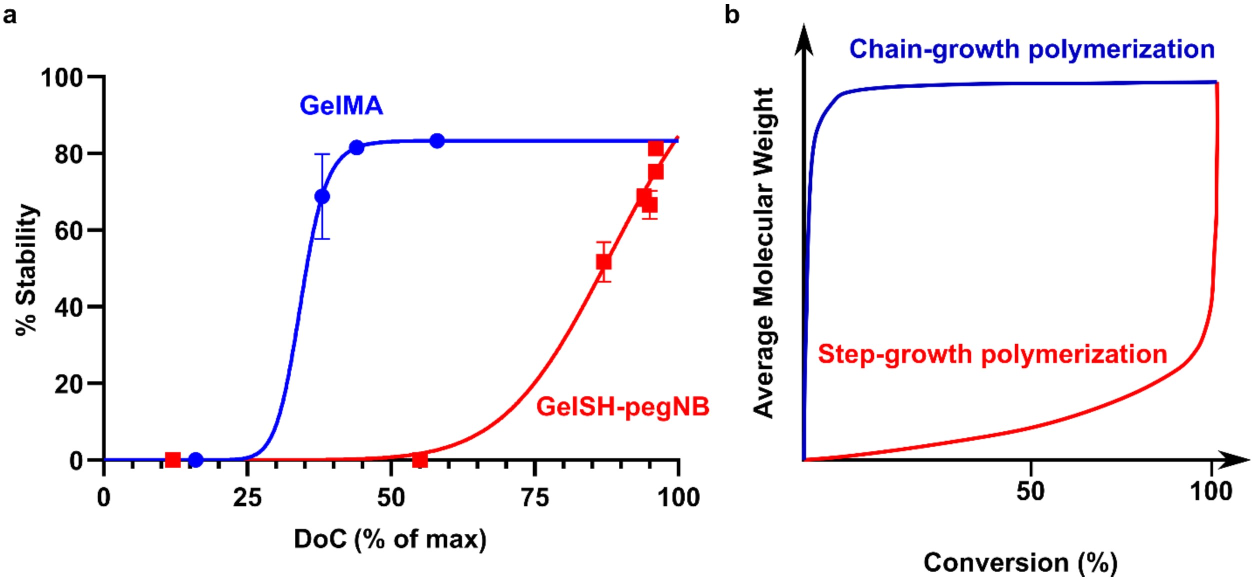


**Supporting Figure S2. The stability of GelMA and GelSH-pegNB trends with DoC in a manner that is analogous to polymer theory.** (a) Overlaid % $DoC_{max}$ vs % stability plots (from Fig 3f and Fig 4g) from photopatterning experiments. GelMA crosslinked through a chain-growth mechanism and exhibited higher stability at relatively earlier DoC, whereas gelSH-pegNB crosslinked by a step-growth mechanism, and did not achieve stability until relatively higher DoC was attained. (b) Classic polymer theory dictates that chain-growth polymers should reach a higher relative molecular weight after fewer crosslinks, whereas step-growth polymers require more crosslinking to reach a maximum weight.[1,2] We posit that the stability of patterned hydrogels is related to the extent of the crosslinked network, thus linking the two plots.
